# Multiple variants of the mitochondrial COI DNA barcode region are prevalent in North European sawflies

**DOI:** 10.1101/2025.04.21.649769

**Authors:** Marko Prous, Santtu Urpilainen, Paul DN Hebert, Evgeny Zakharov, Niina Kiljunen, Marko Mutanen

## Abstract

DNA barcoding, the use of standard segments of DNA to assign specimens to a species, has emerged as a major field of biodiversity research over the last 20 years. Large-scale global initiatives are building DNA barcode reference libraries for animals, fungi, and plants, while pipelines are being developed for metabarcoding-based biomonitoring. The effectiveness of these approaches rests on the premise that much less variation exists within species than between them. While exceptions occur, this principle has been demonstrated to apply in the many animal taxa where the barcode region of the COI gene is effective in species discrimination. Sawflies are an exception to this general pattern because DNA barcodes often fail to distinguish congeneric species, an observation which prompted us to search for an explanation. Using high-throughput single-molecule DNA sequencing to recover COI sequences from thousands of sawflies, we found that single individuals often possess multiple, seemingly functional, full-length DNA barcodes – a phenomenon not documented at similar prevalence in any animal taxon. While the evolutionary causes of multiple variants require further investigation, our observation is remarkable as it violates the one-barcode-one-specimen assumption. The presence of multiple variants of barcodes within individuals does not jeopardize the concept, but its occurrence does introduce a complexity for species inventories based on metabarcoding. They will overestimate the species count when barcode-based operational species units are used as species proxies. Similarly, reference libraries must consider how best to deal with the high frequency of multiple variants in sawflies and any other groups of organisms.

**Significance Statement:** DNA barcoding is revolutionizing biodiversity science by enabling the accurate identification of organisms, accelerating taxonomic workflows, and permitting DNA-based biomonitoring. The DNA barcode region for the animal kingdom, mitochondrial COI, is highly effective in discriminating species in almost all studied animal groups. However, the use DNA barcoding is sometimes complicated by the presence of nuclear pseudogenes (NUMTs) or by variants of the mitogenome itself (heteroplasmy) within individuals. By using high-throughput sequencing (HTS) to analyze thou-sands of specimens, we demonstrate that multiple, seemingly functional, full-length variants of the COI barcode region are frequent in North European sawflies. Since these variants are sometimes very divergent, it is important to consider the impact of this within-individual variability on studies based on DNA barcodes.

## Introduction

Since its introduction 20 years ago (1), DNA barcoding, the use of short, standard fragments of DNA to assign biological specimens to a species, has transformed many areas of biodiversity research (2–4). Due to its effectiveness in distinguishing cryptic taxa, DNA barcoding has facilitated taxonomic research (5, 6). Similarly, it has made it possible to probe species interactions with unprecedented detail and accuracy (7–9) and has enabled efficient mapping of community diversity (10–12). DNA barcoding can accelerate species discovery in ‘dark taxa’, i.e., poorly studied hyperdiverse groups of organisms (13). Besides biological research, the approach is valuable in applied contexts such as forensics (14), biomonitoring (15, 16), detection of invasive species (17), and deterring marketplace fraud (18, 19). Innovative applications based on the concept are constantly emerging, such as the emergent field of metabarcoding (20). Consequently, massive enterprises such as BIOSCAN (https://ibol.org/programs/bioscan/), BGE (https://biodiversitygenomics.eu/), and many national initiatives are presently building DNA barcode reference databases at local, regional or global scales.

In animals, the standard DNA barcode, a 658 bp segment of the mitochondrial COI (cytochrome c oxidase) gene (1), was selected for several reasons. The mitogenome is haploid and shows little or no recombination (21), aiding data interpretation. Multiple copies of COI are present in each cell, facilitating its PCR amplification even from very small organisms. In addition, the evolution of COI is usually rapid enough to discriminate closely related species (22–25). However, DNA barcodes are less effective in distinguishing species in some animal lineages and sawflies (Hymenoptera, Symphyta) are one of them. This compromised performance has been linked to mitonuclear discordance (26–29) and to low barcode divergence among species (30).

The interpretation of DNA barcode data is sometimes complicated by NUMTs, non-coding copies of COI located in nuclear genome. When such NUMTs are long enough to possess the binding sites for both primers used in PCR, they are co-amplified with mtCOI. Because NUMTs are generally less than 300 bp in length, the risk of exposure is greatest when short fragments of COI are amplified so they pose the greatest risk for eDNA or metabarcoding studies (31). While relatively few NUMTs span the barcode region, they do occur (32). As NUMTs are not exposed to natural selection, they often accumulate frameshift mutations and stop codons which allow their discrimination from mtCOI, but long NUMTs without these diagnostic features do occur (32).

Heteroplasmy, the phenomenon where an individual harbours more than one mtDNA haplotype, represents a second potential cause of interpretational complexity. While most organisms inherit a single mitogenome, exceptions have been documented in diverse taxa (33), including plants (34), bivalve molluscs (35), lobsters (36), ants (37), beetles (38, 39), and potentially sawflies (29, 30). Divergent heteroplasmy (>1% divergence) based on whole mitochondrial genomes has been reported in the tuatara (40) and a beetle (39). Heteroplasmy is generally considered uncommon so it is not thought to significantly compromise eDNA or metabarcoding analyses, but large-scale studies on heteroplasmy have not been conducted. An important feature of heteroplasmy is that documenting its presence by Sanger sequencing, until recently the standard method for barcode reference library construction, is difficult. This barrier reflect the fact that heteroplasmy is easily mis-interpreted as contamination in Sanger sequencing chromatograms as both are revealed by the presence of multiple peaks.

The sawfly fauna of northern Europe has been under intensive taxonomic reassessment for the last decade (27, 41–43). As one component of this work, over 20,000 specimens have been DNA barcoded. This analysis revealed unusual patterns of barcode variation. First, barcode sharing was prevalent among closely related species and, second, many individuals of some species possessed two or more barcodes with deep divergence, many of which were examined for nuclear markers to evaluate possible cryptic diversity. The shift from Sanger sequencing to Sequel revealed an unexpected pattern - deep intraspecific variants were not only frequent within species, but also within individuals, suggesting that long NUMTs and/or heteroplasmy were prevalent in sawflies.

In this study, we investigate the frequency of sawflies which possess two or more COI variants by examining 6,763 specimens across 88 genera of sawflies from North Europe, chiefly Finland. Our goals were (1) to ascertain if within-individual variability results from the co-amplification of NUMTs spanning the barcode region which could be recognized because of their possession of diagnostic sequence features (frame shifts, stop codons) or as the co-occurrence of multiple seemingly functional mitochondrial variants within individuals and (2) to assess the prevalence of these full-length within-individual barcode variants among sawflies. As such, our study is the first to investigate the prevalence of long within-individual barcode variants based on screening a massive data set generated using HTS.

## Results

### Characterization of the sawfly dataset

The contamination recognition step removed 2,702 sequences which derived from 1,939 specimens (Table 1). The removal of diagnosable NUMTs excised 3,458 sequences (1,768 specimens). The remaining dataset contained 7,879 sequence variants derived from 6,173 specimens. When excluding variants with < 3 reads, the dataset contained 5,474 specimens with 6,064 sequence variants.

**Table 1.**
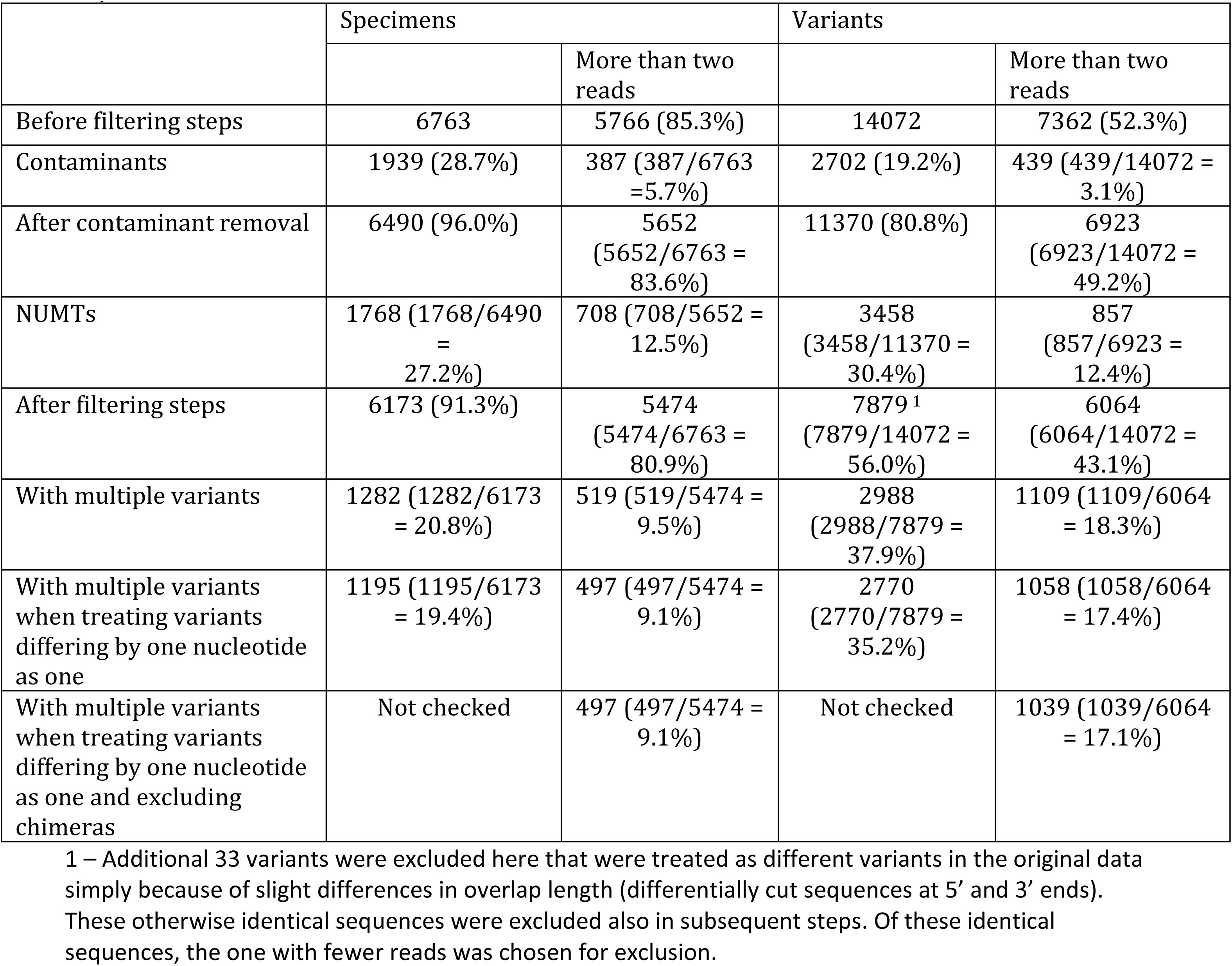
Number of specimens and COI sequence variants before and after consecutive filtering steps.

After exclusion of contaminants and diagnosable NUMTs, 20.8% of specimens contained two or more COI variants (Table 1). After excluding sequences with fewer than three reads, 9.5% of specimens showed two or more variants. When only retaining variants differing by more than one nucleotide, 19.4% (all sequences) or 9.1% (sequences supported by 3 or more reads) of the specimens had more than one variant (Table 1). Finally, the removal of chimeras in the dataset with sequences supported by more than two reads left 497 specimens (9.1%) with 1,039 variants (Table 1, Supplementary Fig. 1).

### Co-presence of intraindividual variants in sawflies

After exclusion of diagnosable NUMTs, a fifth (20.8%) of the specimens (1,282/6,173) in the sawfly dataset possessed intraindividual variants. Most had two variants, but as many as 10 were observed. The median read count for the major variant in these specimens was 18 (maximum = 126) while the median read count for the minor variant was 2 (maximum = 35). When present, the other variants had a median read count of 1 and maximum of 15.

Once sequences with fewer than three reads were excluded (Table 1), 9.5% of the specimens (519/5,474) in the sawfly dataset retained intraindividual variants. Most of these specimens had two variants, with a maximum of four. For the major variant, the median read count was 19 with a maximum of 110. For the next commonest variant, the median read count was 5 with maximum of 33. Minor variants had a median read count of 4 and a maximum of 15.

Considering only the dataset based on sequences supported by at least three reads, the following genera with at least 8 specimens showed a high incidence (>30%) of intraindividual variation (Supplementary Table 2): *Fenella* (100%), *Monophadnoides* (80%), *Eriocampa* (69%), *Nesoselandria* (52%), *Cladius* (36%), *Cephalcia* (36%), *Nematus* (33%), and *Xiphydria* (33%).

Considering all 347 species or species groups, 110 included specimens with more than one intraindividual variant (dataset with sequences supported by at least three reads). Considering only species or species-groups represented by at least 11 specimens, 16 showed more than 30% of their members with two or more COI variants. They included *Ametastegia tenera* (92%), *Dolerus subarcticus* (82%), *Monophadnoides rubi* (80%), *Euura obducta* (79%), *Nesoselandria morio* (52%), *Cladius pectinicornis* (45%), *Dolerus vestigialis* (41%), *Empria pallimacula* (37%), *Cephalcia* spp (36%), *Claremontia brevicornis* (35%), *Cladius brullei* (33%), *Dineura virididorsata* (32%), *Cladius compressicornis* (32%), *Euura vaga* (32%), *Euura myosotidis* (31%), and *Tenthredo atra* group (30%). Supplementary Tables 1–2 provide details on all species and genera with multiple intraindividual variants.

When specimens with multiple apparently functional variants were compared to specimens with a single variant, there were no statistically significant differences in sex ratios (p-values 0.11-0.76 Table 2).

**Table 2.**
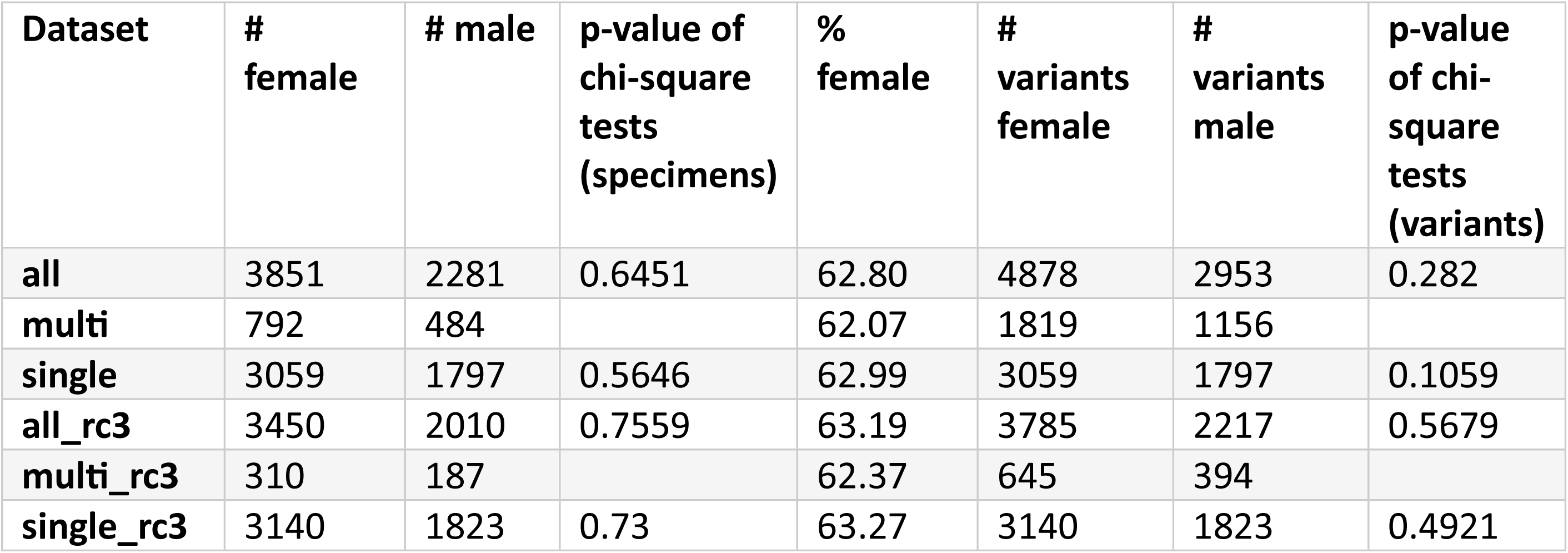
Number of female and male specimens (excluding larvae) and variants in different datasets. “all” includes specimens both with single and multiple apparently functional intraindividual COI variants. “multi” dataset contains only specimens with multiple intraindividual variants and “single” contains only specimens with one variant. “rc3” means that only sequences supported by at least three reads were considered. Possible PCR chimeras were excluded and variants differing by single nucleotide were treated as single variant in rc3 datasets. Chi-square tests are comparisons of male and female numbers between multiple variant and all or single datasets.

### Intraindividual divergences of sawflies

The maximum likelihood tree for the most stringently filtered dataset of specimens with intraindividual variants is provided as supplementary material, but some remarkable examples are illustrated in Figs 1–4. Specimens of *Empria pumila* possessed up to four variants with divergences from 2.3 – 5.7% while two variants with up to 8.6% divergence were detected in *E. pallimacula* (Fig. 1). Two closely related species of *Tenthredo* (*T. ferruginea*, *T. silensis*) possessed intraindividual variants showing 4.8 - 6.9% divergence (Fig. 2). Specimens of *Cephalcia* (number of species uncertain) formed two main barcode lineages which are not shared by specimens (except ZMUO.033841), but those within each group are polyphyletic. The maximum genetic distances between intraindividual variants within these groups are 4.8% or 6.6% (Fig. 3). There were also two groups within *Cladius pectinicornis*, which do not seem to share specimens (Fig. 4). Within the groups, the maximum distances between intraindividual variants are 3.4% and 3.8%.

**Figure 1.**
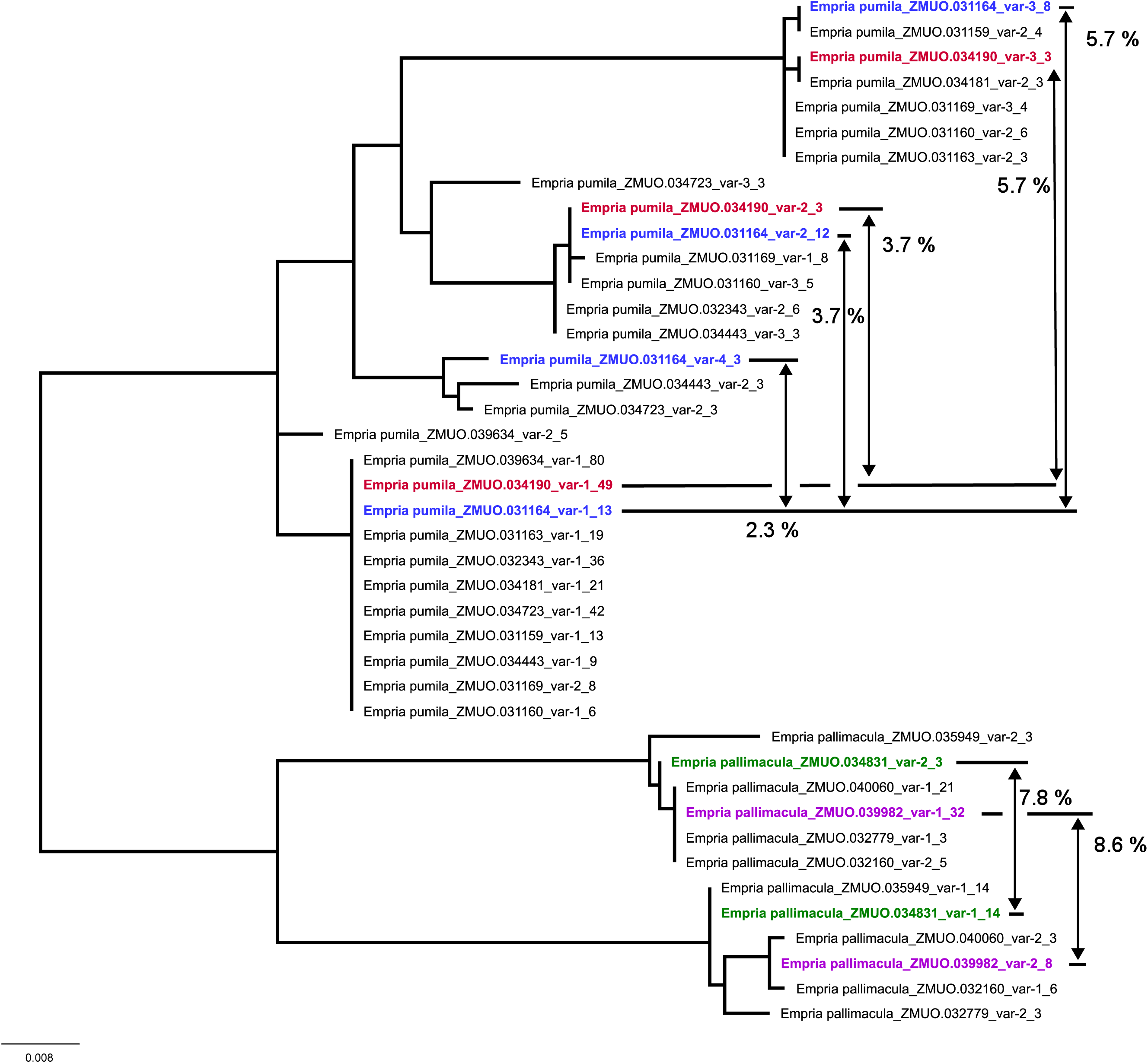
Specimens of *Empria pallimacula* and *E. pumila* with seemingly functional intraindividual COI variants. p-distances between the variants are shown for selected individuals.

**Figure 2.**
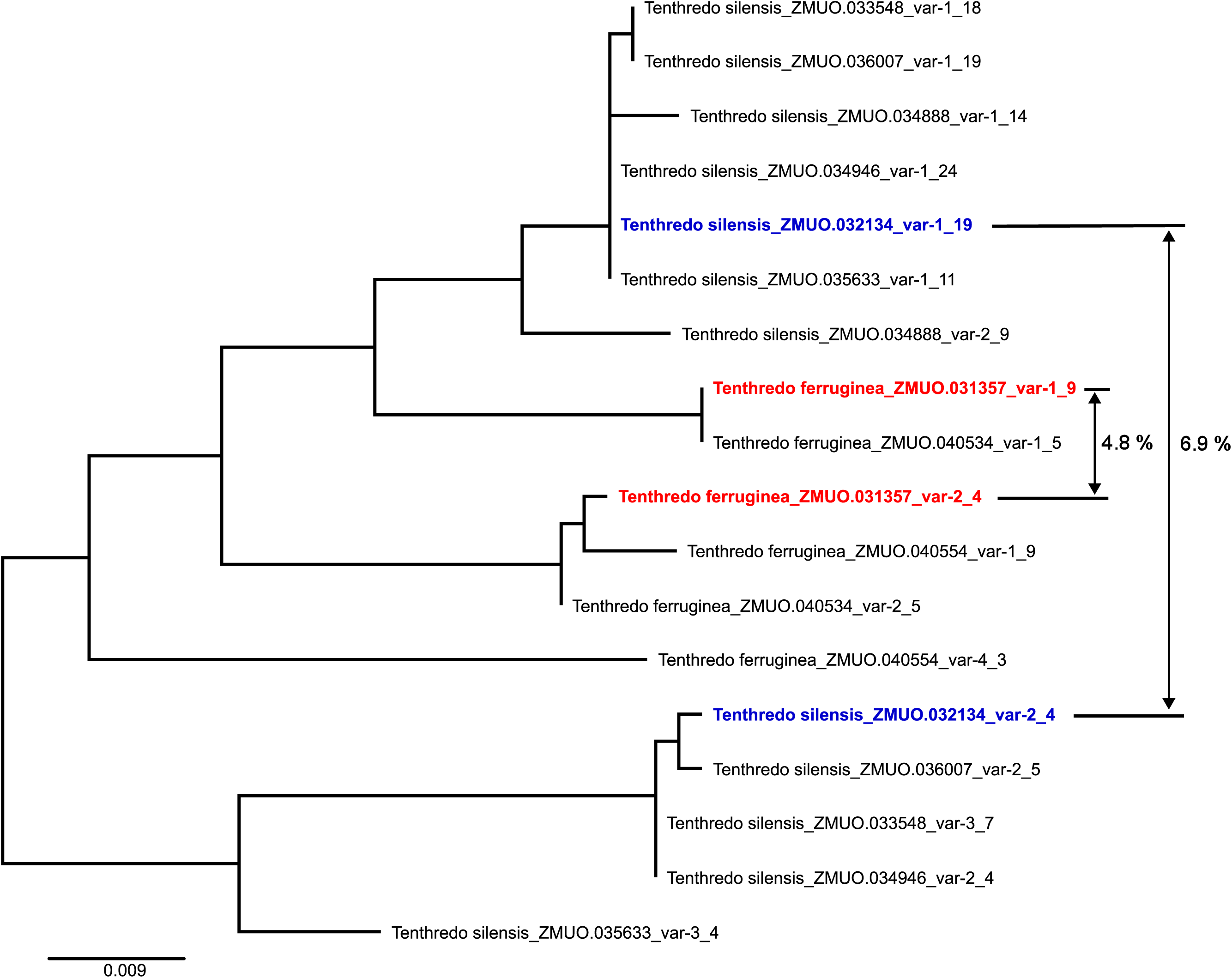
Specimens of *Tenthredo ferruginea* and *T. silensis* with seemingly functional intraindividual COI variants. p-distances between the variants are shown for selected individuals.

**Figure 3.**
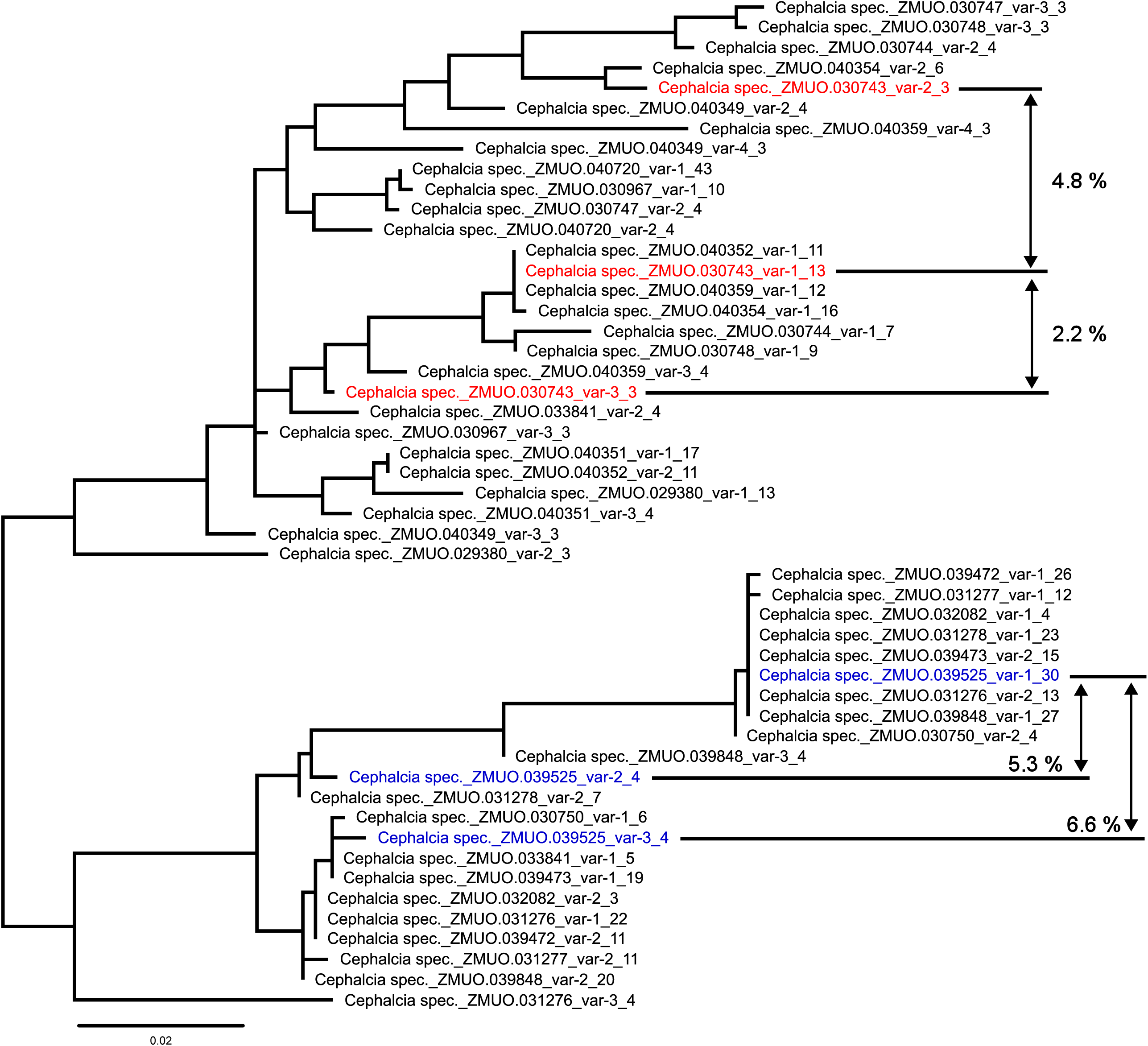
Specimens of *Cephalcia* (number of species uncertain) with seemingly functional intraindividual COI variants. p-distances between the variants are shown for selected individuals.

**Figure 4.**
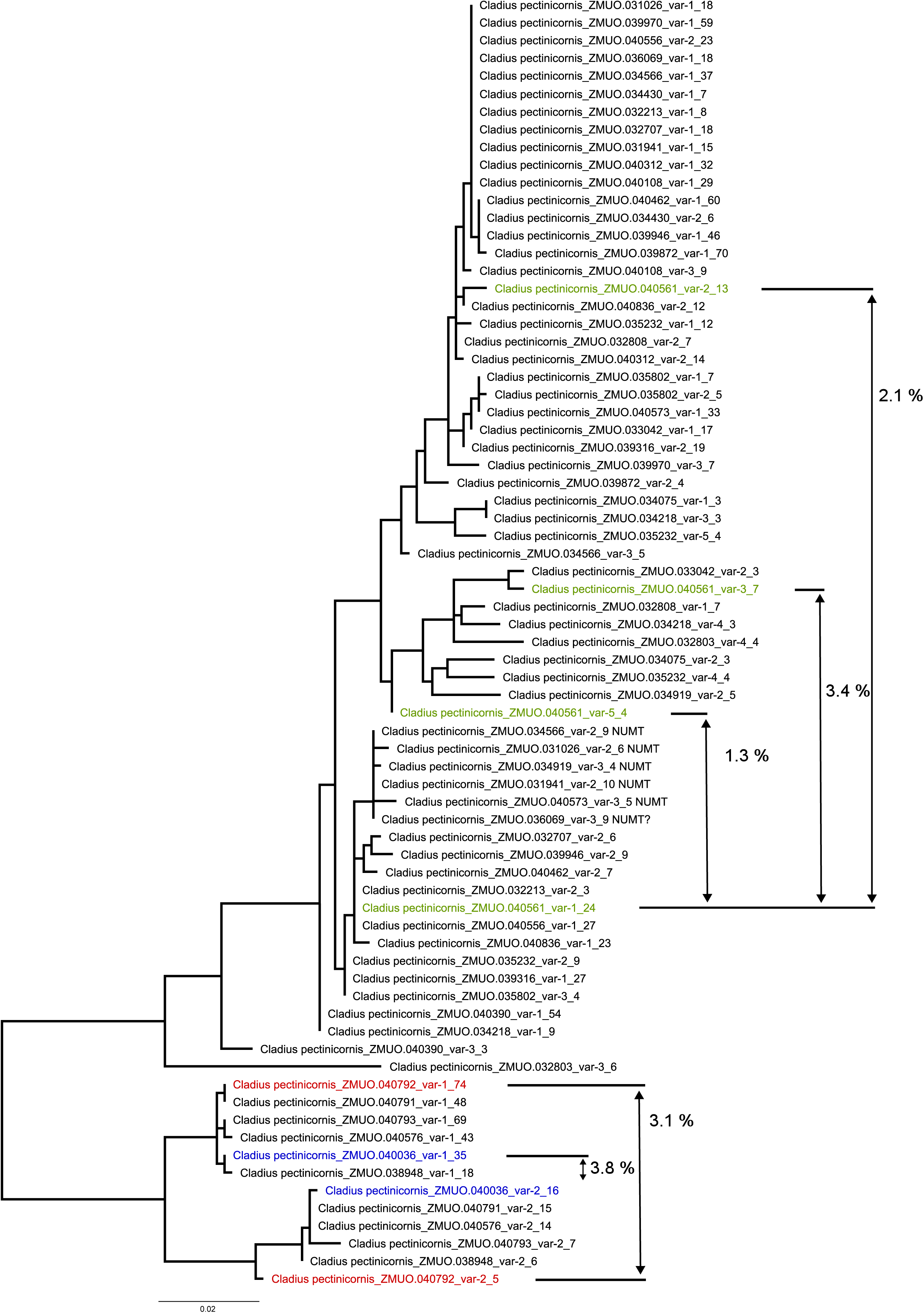
Specimens of *Cladius pectinicornis* with seemingly functional intraindividual COI variants. p-distances between the variants are shown for selected individuals.

## Discussion

With 5,474 specimens and about 347 species or species groups of sawflies represented in the final validated dataset, our study represents the first large-scale investigation of intraindividual full-length mtCOI DNA barcode variants. Our observations might potentially arise from sequencing errors or overlooked contaminants. However, sequencing errors were eliminated by excluding reads that were only recovered once (although Sequel reads have high fidelity). Bioinformatics pipelines do provide an efficient way to recognize sequence variation derived from contamination, particularly when the taxa in a sample is known and when comprehensive reference libraries are available. Given the prevalence of intraindividual long variants, our observations are important as they impact results gathered through metabarcoding or eDNA analysis because such intra-individual variants inflate the estimated species number. Variation of this type is certainly not restricted to sawflies, but its prevalence varies among taxonomic groups. For example, NUMTs are more prevalent in insect species with direct development or incomplete metamorphosis than in those employing complete metamorphosis, reflecting genome size differences (32). Elucidating the distribution of intraindividual variants across all taxa is an obvious next step. The emergence of high-throughput DNA sequencers enables their efficient detection as demonstrated here but may also provide avenues to mitigate its downsides.

Our analysis demonstrates that HTS produces many sequences that seemingly represent the target region but are actually NUMTs or heteroplasmy. NUMTs are a well-recognized complication for DNA barcoding, and recognition of those with frameshift mutations or stop codons is straightforward However, the diagnosis of shorter NUMTs if often a challenging. A recent analysis of 1,002 insect genomes revealed that NUMTs are ubiquitous in insects (32). While it found that most NUMTs are short and that that most NUMTs spanning the barcode region possess indels or premature stop codons, this was not always the case. Some long NUMTS lack diagnostic features, but they were uncommon (32). By contrast, our dataset includes thousands of variants that remain unrecognizable as NUMTs, provided that the intraindividual variants really represent NUMTs. However, at least in the species rich genus Euura (25% of specimens in our dataset), 25% of about 1000 specimens show seemingly functional intraindividual variants even based on 1078–1087 bp COI amplicons (43). Hence the key questions are: a) Why are long intraindividual variants with no indels or stop codons so prevalent in sawflies, and b) in addition to NUMTs, might processes such as heteroplasmy explain their occurrence?

Some of the intraindividual variants which appear fully functional possess more than 2% sequence divergence, a threshold often used to flag putative cryptic species. Moreover, cases of intraindividual variants were frequent in many genera with the same or similar variants occurring in different species, compromising the ability of DNA barcodes to separate them. Further, the observed incidence of intraindividual variability is likely an underestimate because the mitochondrial variants varied in frequency among individuals. Therefore, any single high-throughput run is unlikely to recover all variants present in a species.

In Sanger sequencing, the presence of two or more co-amplified barcode variants generates double peaks in chromatograms, but there is no way to ascertain if they reflect contamination, heteroplasmy, or NUMTs. Deeper insights into these situations only became possible with the emergence of single molecule sequencers capable of generating long reads (e.g., PacBio, Oxford Nanopore). Although the detection of co-amplified variants is now efficient, it remains difficult to discriminate heteroplasmic variants from long NUMTs which lack stop codons or frameshift mutations. The discrimination of such cases requires either transcriptomic analysis because NUMTs are not transcribed while heteroplasmic variants are, or whole genome sequencing to localize NUMTs in the nuclear genome.

Can we exclude the possibility that the cases of deep divergence between multiple variants reflect heteroplasmy? Past studies have shown that most cases of heteroplasmy involve very low levels of divergence (56, 57). The seemingly functional intraindividual COI variants of ∼650 bp barcode using PacBio are also found with Nanopore sequencing of ∼1080 bp mtCOI fragment (43). This observation supports the heteroplasmy hypothesis, although Hebert et al (32) observed a few dozen NUMTs spanning over 1,500 bp but lacking frameshift mutations or stop codons. As such, the detection of long variants in sawflies is not compelling evidence for the heteroplasmy hypothesis. It has been suggested that paternal leakage of mitochondria increases with increasing genetic divergence between the parents (e.g. in case of interspecific hybrids) due to failure of paternal mtDNA elimination (33, 58). For example, divergent parent mitochondrial types are frequently detected in *Pelophylax* frogs but only in hybrids (59). In sawflies, mitonuclear discordance is common and may reflect mitochondrial introgression (26, 28), which would be consistent with the prevalence of possible heteroplasmy detected in this study.

If the “functional” intraindividual variants are NUMTs, they would be expected to be more prevalent in diploid females than haploid males if the NUMTs are polymorphic. However, no difference in sex ratio were found between specimens with and without multiple variants (Table 2), a result that can be explained if the NUMTs are not polymorphic.

To rule out the NUMT hypothesis, complete nuclear and mitochondrial genomes should be sequenced. For example, the chromosome level genome assembly for the sawfly *Tenthredo mesomela* (https://www.ncbi.nlm.nih.gov/datasets/genome/GCA_943736025.1/), includes two nuclear scaffolds (OX031020, OX031013) with ∼7000 bp segments of the mitogenome (GenBank accession OX031023) with sequence identity of 98–99%. Another possibility to distinguish between heteroplasmy and NUMTs would be to screen for the multiple variants in different specimens and populations of the same species (39). Divergent NUMTs should be present in all specimens, while heteroplasmic variants should occur in only some of individuals (39).

As long intraindividual barcode variants are widespread in sawflies: it is important to consider how best to mitigate their effect on both the identification of single specimens and species inferences from metabarcoding. We recognize two concrete actions towards this goal. First, DNA barcoding should transition to single molecule sequencing instead of Sanger, because only the former can separate the multiple variants. The shift from Sanger to high-throughput sequencing is underway and, before long, all barcoding will be based on the latter approach as it is much less expensive. Second, as long as barcoding applications include PCR step, intraindividual barcode variants will be co-amplified. The much higher copy number of mtCOI should reduce exposure to NUMTs (32), but in our experience with sawflies, any of these variants could end up to BOLD as the employed bioinformatic pipelines selects, by default, the variant with most reads to represent the specimen’s barcode. As such, variants persist in data sets, a practical solution could be to index them as alternative barcodes. Co-amplified, seemingly functional variants should become an integral part of the reference libraries as otherwise such ‘ghost’ species will be retained in metagenomics data sets. Alternatively, DNA barcoding could be based on transcribed DNA which would enable the exclusion of non-transcribed NUMTs, but this is only possible for fresh material as RNA degrades quickly.

In conclusion, analysis of 6,000 individuals using high-throughput single-molecule sequencing platforms frequently recovered multiple long mtCOI variants from single specimens of sawflies. While the factors underlying this phenomenon require further investigation, their presence has significant implications for DNA barcoding and metabarcoding studies on this group. While the documented observations form a complication for such ambitious enterprises, single molecule-based sequencing platforms also will provide efficient ways to overcome them. Barcode library construction initiatives therefore should increasingly turn using high-throughput platforms.

## Materials and Methods

Our initial datasets were recovered using circular consensus sequences (CCSs) of mtCOI amplicons generated of Sequel /Sequel II platforms at the Centre for Biodiversity Genomics (CBG). This analytical path generates a read count for every CCS recovered from a specimen, a taxonomic assignment based on the reference library in BOLD, and a percent similarity to the closest reference sequence. This dataset, which was derived from the analysis of 6,773 specimens representing about 480 species included 14,090 sequence variants. After initial filtering (see Supplementary material), 14,072 sequences from 6,763 sawflies were retained for further analyses. Some identifications were based on less than 20 bases of overlap in the mBRAVE software which led to incorrect taxonomic assignments. As a result, the dataset (14,072 sequences) was re-identified against the reference sawfly dataset (see below). The main analytical steps were 1) exclusion of sequences reflecting cross-contamination or non-target sequences (e.g., endosymbionts, parasitoids), 2), exclusion of NUMTs with diagnostic features, and 3) identification of the remaining specimens with multiple variants. Cases of contamination were defined as those with sequences matching specimens from a different genus. Diagnosable NUMTs were defined as sequences with stop codons and/or frameshift indels (1 or 2 bp insertions/deletions). We provide counts for each of these three categories of sequences (contaminants, diagnosable NUMTs, potentially functional sequences) based on either all CSSs or restricted to CSSs with at least three reads to minimize exposure to sequencing error.

The sawfly reference dataset contains published and unpublished mtCOI sequences connected to the Electronic World Catalog of Symphyta (44). Due to the lack of consistency in taxonomic names across databases (most linked to the recent generic revision of the Nematinae (45)) as well as gaps or errors in identification, the names of specimens were corrected to a genus or species level before further analyses following the latest taxonomy (43, 44).

Sequences derived from contamination was removed based on four filters. Sequences were first excluded if they were proteobacterial or derived from a different family than the source specimen. After this filtration, the remaining sequences were re-identified based on a reference dataset using BlastN (46) with an e-value threshold of 1E-150, and excluded if the assigned genus did not match the generic assignment based on morphology. 4) Some additional cases of contamination were detected within genera, involving distantly related species or species-groups not known to share barcodes.

NUMT detection was done in three steps 1) detection of frameshifts with BlastN (46) against a nucleotide reference dataset, 2) detection of stop codons with BlastX (46) against an amino acid reference dataset, and 3) manual checks of sequences for which the Blast results were ambiguous regarding frameshifts. Reads longer than the amplicon region (e.g., >659 bp) were retained if stop codons or frameshifts were only detected in the primer regions. Reads significantly shorter than expected amplicon length (e.g., <212 amino acids, a 658 bp amplicon should include 219 aa) were checked to ascertain if they were obviously chimeric and those detected were excluded. Although sequences with more than three aa deletions are probably NUMTs, they were retained for consistency (there were only five sequences with more than two aa deletions). The remaining (11,370) sequences were 1) aligned with MAFFT (47) at a nucleotide level to establish a common translation frame, 2) translated to protein (using the invertebrate genetic code), 3) aligned again with MAFFT at an amino acid (aa) level, and 4) using pal2nal (48) to recover a the nucleotide alignment based on the aa alignment. This enabled the detection of additional problematic sequences (overlooked stop codons close to the ends of sequences, frameshift and unusually long indels). See supplementary material for additional details.

As an additional quality control, we also reported the number of possible NUMTs/heteroplasmic variants by only considering sequences differing by more than one bp substitution or indel. Although this criterion will undoubtedly exclude many true variants, those remaining are more reliable, particularly when supported by multiple reads.

Finally, the more stringently filtered dataset (CSSs supported by >2 reads and excluding intraindividual variants differing by 1 nucleotide) containing specimens with seemingly functional intraindividual COI variants was manually examined for PCR chimeras.

A phylogenetic tree was generated for the most stringently filtered dataset with FastTree (49). Prior to phylogenetic analyses, the primer sequences were trimmed.

Analyses were done using custom scripts written in R (50) using packages plyr (51), dplyr (52), xlsx (53) and tidyr (54).

For selected species or species groups with multiple mitochondrial variants, p-distances were calculated in R using ape (55). A chi-square test in R (50) was used to check if the sex ratio differed between specimens with multiple apparently functional intraindividual COI variants and those with single COI variant.

## Acknowledgments

This study has been supported in part by the Finnish Research Council through research infrastructure funding to FinBIF consortium and in part by the New Frontiers in Research Fund and by the Canada Foundation for Innovation’s Major Science Infrastructures program. MP is supported by the Estonian Research Council grant STP42. We also want to remember our co-author and PhD student Santtu Urpilainen who sadly passed away due to illness before the publication of this work.

## Supporting Information for

### Supporting Information Text

#### Filtering methods

##### Preliminary filtering steps

The initial sawfly dataset contains 6773 specimens with 14,090 sequence variants. 17 sequences (9 specimens) were excluded due to processing error (before samples were sent for sequencing) that caused a specimen and sequence mismatch. In addition, one specimen of Apocrita (Eurydinota GN.148) was removed. After their exclusion, 14,072 sequence variants and 6763 specimens remained (DatasetS1.xlsx).

##### Exclusion of clear (cross-)contaminations or other non-target sequences (endosymbionts, parasitoids etc.)

Due to a bug in the mBRAVE software (identifications based on very short overlap, less than 20 bases) sequences were re-identified against a sawfly reference database. Blast-database was constructed using reference file DatasetS2.fas.

The workflow was as follows. Sequence variants initially identified as Proteobacteria and sequences that did not match the expected family identification were treated as contaminants. The remaining sequences were re-identified using a BLAST search with e-value of 1E-150. After re-identification, sequence variants that did not match the expected genus were considered also contaminants. Some additional contaminants were detected within genera, involving distantly related species or species-groups not known to share barcodes. The remaining ones were exported to a fasta-file for NUMT detection. All sequences identified as contaminations are listed in DatasetS1.xlsx.

##### Exclusion of possible NUMTs

We define diagnosable NUMTs as sequences containing stop codons or frame shifting indels (insertions or deletions). Detection of NUMTs was done in three steps.

i. After removing contaminants identified in the previous step, the remaining sequences were subjected to BlastN (1) search against a nucleotide reference dataset (DatasetS2.fas) with following settings, -evalue 1E-150 -max_target_seqs 100. After BlastN, sequences without indels and those with the sum of gaps (indels) divisible by three, were retained.
ii. Sequences left from the previous step were subjected to BlastX (1) search against an amino acid reference database (DatasetS3.fas) with settings -evalue 1E-50 -max_target_seqs 200. After BlastX, sequences with stop codons (notated by an asterisk ‘*’) were excluded.
iii. Three different manual checks were done for the sequences that were left ambiguous regarding frameshift insertions or deletions. First, sequences with ambiguous gaps from nucleotide blast-search from step i) were filtered out if they contained stop codons based on results from step ii). The remaining sequences were aligned against their closest reference using MAFFT (2). After alignment the sequences were manually checked for frameshifting insertions and deletions. Sequences with frameshifting indels were removed.

For additional quality control, sequences longer than the amplicon region (>659 bp) were checked for stop codons and indels in the primer regions. Sequences longer than >659 bp and containing indels or stop codons were combined with their closest match and aligned using MAFFT (2). Primer regions were delimited based on standard barcoding primers (LepF1/LepR1). Sequences with indels only in the primer regions were recovered for subsequent analyses. Based on BlastX search (same settings as above) no stop codons were detected in these sequences.

Sequences that were shorter than the expected amplicon length (<212 aa) and that did not contain indels or stop codons based on earlier steps were checked manually for ambiguous and problematic sequences (e.g. chimeric) and aligned against their closest reference using MAFFT. Sequences deemed chimeric were excluded.

The remaining sequences were aligned with MAFFT to detect additional problematic sequences. To improve the alignment, the initial alignment was split into two. The first non-problematic part (only varying degrees of missing data in different specimens) until the alignment position without any missing data and corresponding to a third codon position was saved as separate alignment. The second alignment part was saved by removing all gaps, translated into protein using codon table 5 (invertebrate genetic code), and aligned again with MAFFT. The resulting protein alignment was used to get corresponding and more reliable nucleotide alignment with pal2nal (3). Two parts of the alignment were then merged. Examination revealed a few additional sequences with frameshifts or unusually long indels which were removed. Removed were also about 30 additional sequences with stop codons very close to the ends of the sequence that were not detected in previous analyses. All sequences identified as NUMTs, although some of them may be sequencing errors, are listed in DatasetS1.xlsx.

##### Identification of remaining specimens with multiple variants

Before analysis, additional taxonomic curation at the species level was done for two main reasons: 1) to resolve differences between the reference and the PacBio dataset (e.g., the closest match in the reference dataset was a species missing from the PacBio dataset), 2) identify certain species to species or species-group level because of unresolved or difficult taxonomy or because of known unreliability of COI barcode identification, 3) and correct identifications, e.g. when different variants of the same specimen where identified differently but could not be classified as contaminations. All the results are reported based on all circular consensus sequences (CSSs) or sequences based on at least three CSSs.

As an additional quality control, we also report number of variants by only considering intraindividual CSSs that differ from each other by more than one substitution or indel. Sequences from the same specimen were compared to each other using BlastN. From the blast results (columns “mismatch” and “gaps”), variants that differ from each other by one nucleotide were counted. For sequences without indels (gaps) or substitutions (mismatch) in the blast output, we further checked for the match between the alignment and sequence lengths. Sequences were counted as having one nucleotide difference also when: 1) subject (sequence to be compared) length was equal or longer than query (sequence used for comparison) length and query length was one nucleotide longer than alignment length, or 2) query length was equal or longer than subject length and subject length was one nucleotide longer than alignment length. The intraindividual variants differing from each other by one substitution or indel (including variants classified as NUMTs) are listed in DatasetS1.xlsx. The secondary intraindividual variants that were considered different variants in the original dataset simply because of slightly different overlap length are listed in DatasetS1.xlsx.

The mean, median and maximum number of variants per specimen were calculated for multiple variant datasets (without NUMTs and contaminants). The mean, median and maximum read count was calculated for major, second most common, and minor variants.

Finally, we manually checked for PCR chimeras for the most stringently filtered multiple variants dataset: excluding variants supported by less than three reads and treating variants differing by one nucleotide as one. The within specimen variants were compared to each other taking into account also sequences from the same specimen classified as NUMTs and/or contaminations (if present). PCR chimeras should be more than two times less abundant than parent variants and should not have unique substitutions or indels. Nevertheless, we conservatively treated some ambiguous cases as chimeras (read count same or almost same to one of the parent variants or with one unique substitution, but otherwise look like chimeras) and removed them. Altogether, 22 possibly chimeric sequences were removed. These are listed in DatasetS1.xlsx.

Based on the above chimera filtered dataset, we summarize the taxonomic distribution of specimens with multiple variants at the species and genus level (tables 1–2).

**Fig. S1.**
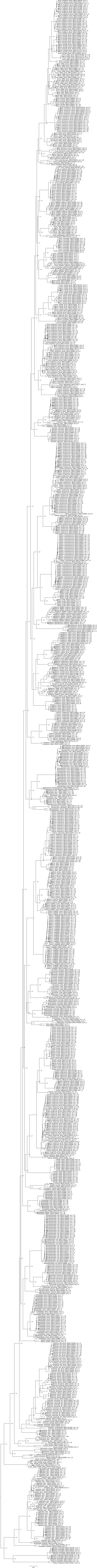
Maximum likelihood tree (FastTree) for the most stringently filtered dataset of specimens with intraindividual variants (sequences supported by at least three reads, with minimum of two differences between the intraindividual variants, no contaminations, no variants identifiable as NUMTs, no chimeras).

**Table S1.**
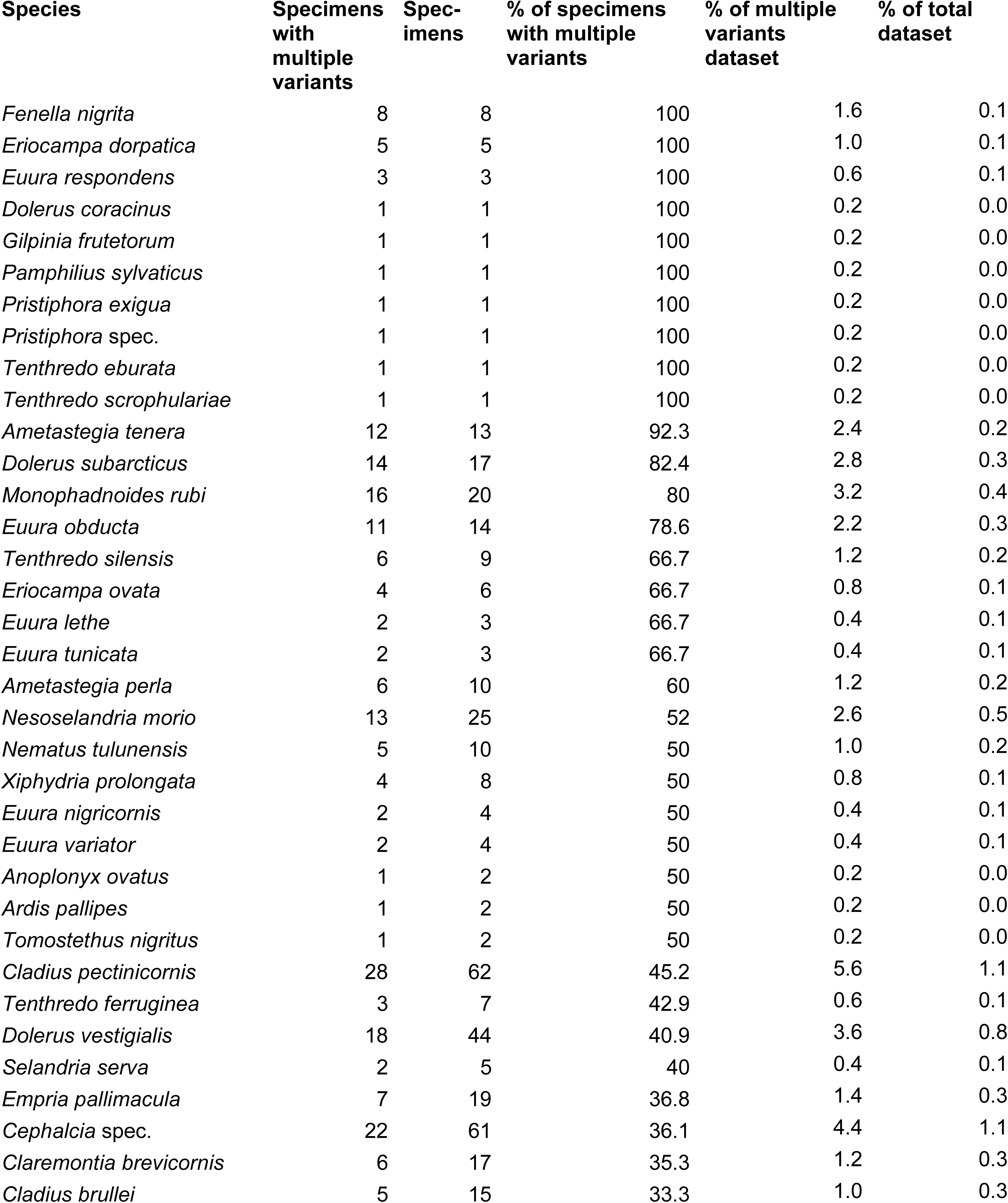
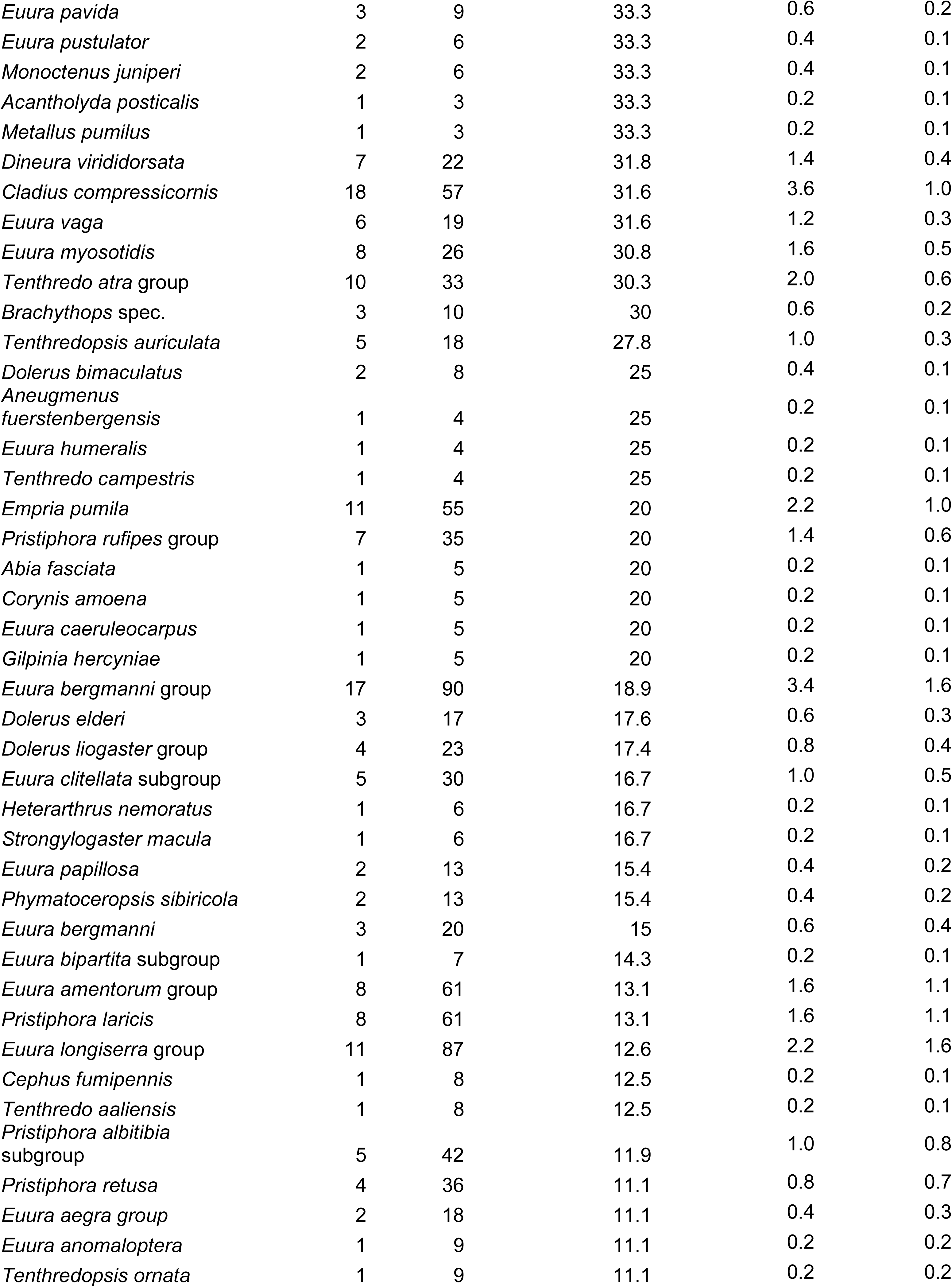
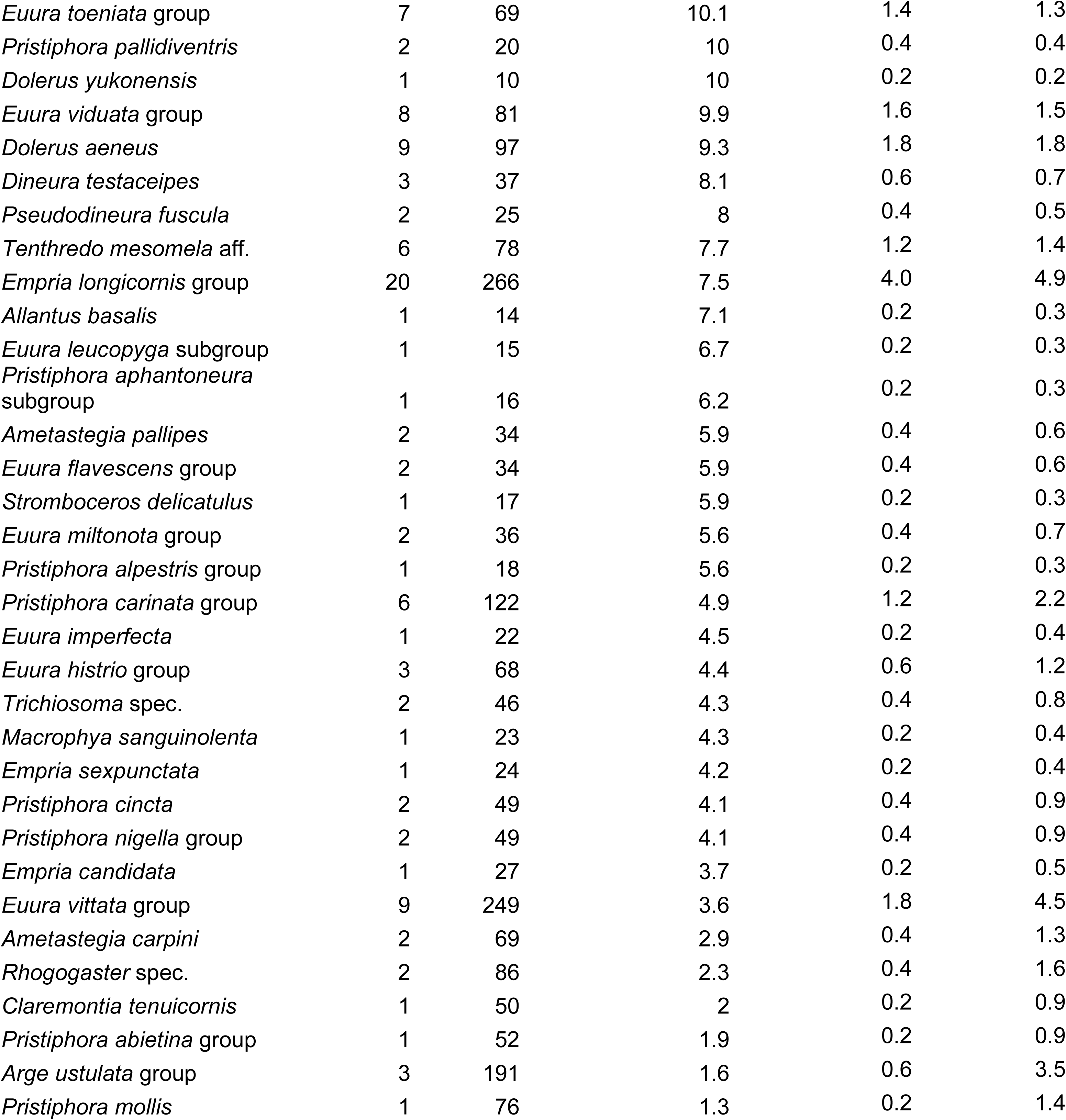
List of species with multiple variants supported by more than two CCS reads and after all filtering steps (contamination, NUMT, chimera removal and excluding variants differing by single nucleotide). The filtered multiple variant dataset contains 497 specimens and total dataset 5474 specimens.

**Table S2.**
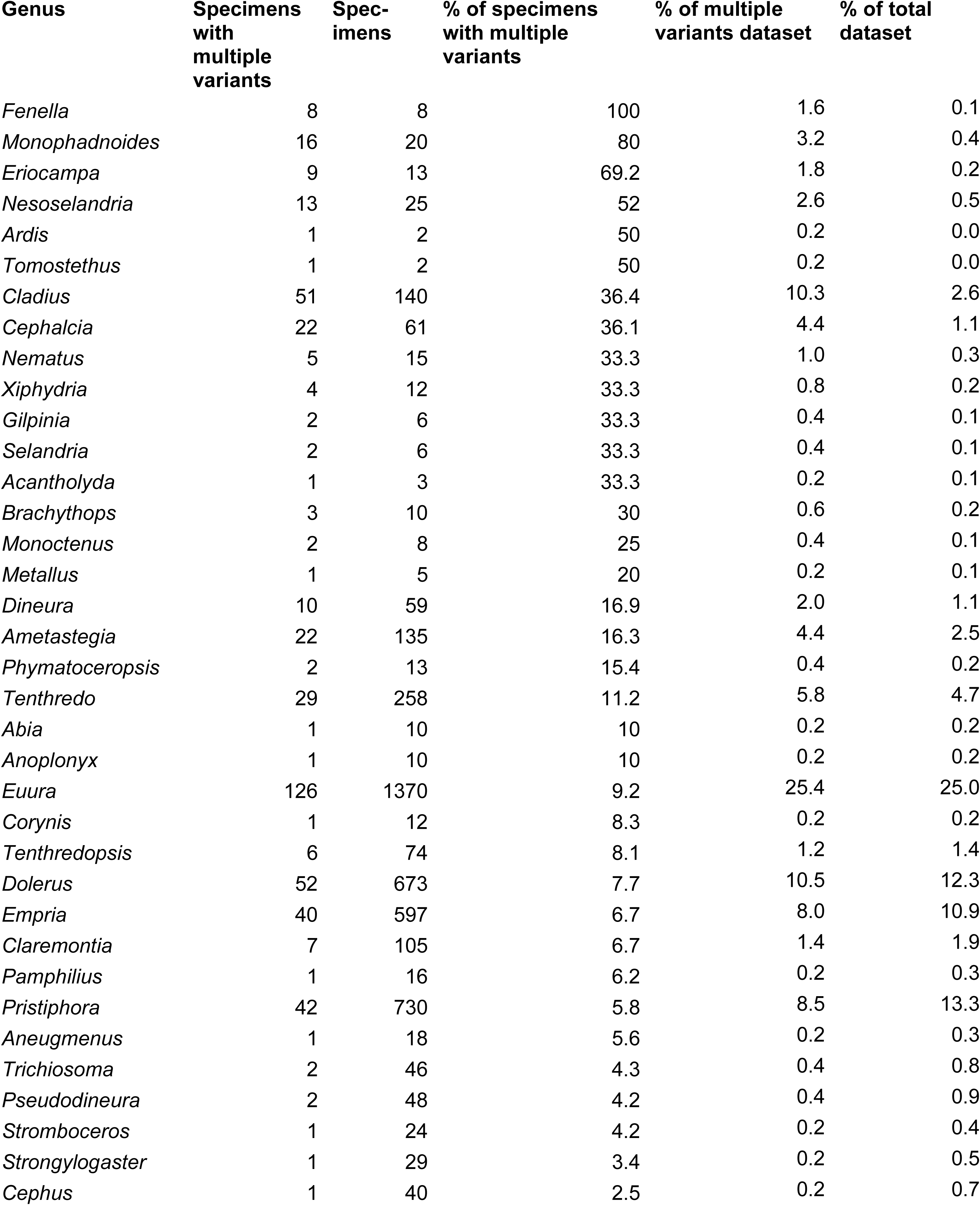
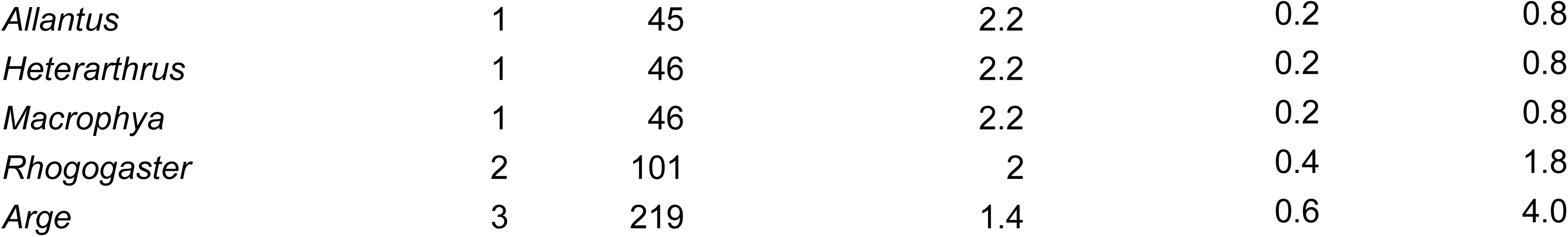
List of genera with multiple variants supported by more than two CCS reads and after all filtering steps (contamination, NUMT, chimera removal and excluding variants differing by single nucleotide). The filtered multiple variant dataset contains 497 specimens and total dataset 5474 specimens.

**Dataset S1 (separate file).** Data tables containing original sawfly PacBio CCS reads, variants after contamination and NUMT filtering, sequence IDs for variants identified as contaminations, variants identified as NUMTs, variants differing by single nucleotide (substitution or indel), identical variants, variants (supported by at least three reads) identified as PCR chimeras.

**Dataset S2 (separate file).** Reference sawfly COI sequences in fastA format used for re-identification of PacBio reads.

**Dataset S3 (separate file).** Reference sawfly COI protein sequences in fastA used for re-identification of PacBio reads.

**Dataset S4 (separate file).** Sawfly specimens with intraindividual COI variants (fastA alignment used to build the tree in Fig. S1) after the most stringent filtering steps (sequences supported by at least three reads, with minimum of two differences between the intraindividual variants, no contaminations, no variants identifiable as NUMTs, no chimeras).

